# Differential local stability governs the metamorphic fold-switch of bacterial virulence factor RfaH

**DOI:** 10.1101/629477

**Authors:** P. Galaz-Davison, J.A. Molina, S. Silletti, E.A. Komives, S.H. Knauer, I. Artsimovitch, C.A. Ramírez-Sarmiento

## Abstract

A regulatory factor RfaH, present in many Gram-negative bacterial pathogens, is required for transcription and translation of long operons encoding virulence determinants. *Escherichia coli* RfaH action is controlled by a unique large-scale structural rearrangement triggered by recruitment to transcription elongation complexes through a specific DNA sequence within these operons. Upon recruitment, the C-terminal domain of this two-domain protein refolds from an α-hairpin, which is bound to the RNA polymerase binding site within the N-terminal domain of RfaH, into an unbound β-barrel that interacts with the ribosome to enable translation. Although structures of the autoinhibited (α-hairpin) and active (β-barrel) states and plausible refolding pathways have been reported, how this reversible switch is encoded within RfaH sequence and structure is poorly understood. Here, we combined hydrogen-deuterium exchange measurements by mass spectrometry and nuclear magnetic resonance with molecular dynamics to evaluate the differential local stability between both RfaH folds. Deuteron incorporation reveals that the tip of the C-terminal hairpin (residues 125-145) is stably folded in the autoinhibited state (∼20% deuteron incorporation), while the rest of this domain is highly flexible (>40% deuteron incorporation) and its flexibility only decreases in the β-folded state. Computationally-predicted ΔGs agree with these results by displaying similar anisotropic stability within the tip of the α-hairpin and on neighboring N-terminal domain residues. Remarkably, the β-folded state shows comparable stability to non-metamorphic homologs. Our findings provide information critical for understanding the metamorphic behavior of RfaH and other chameleon proteins, and for devising targeted strategies to combat bacterial diseases.

**Significance:** Infections caused by Gram-negative bacteria are a worldwide health threat due to rapid acquisition of antibiotic resistance. RfaH, a protein essential for virulence in several Gram-negative pathogens, undergoes a large-scale structural rearrangement in which one RfaH domain completely refolds. Refolding transforms RfaH from an inactive state that restricts RfaH recruitment to a few target genes into an active state that binds to, and couples, transcription and translation machineries to elicit dramatic activation of gene expression. However, the molecular basis of this unique conformational change is poorly understood. Here, we combine molecular dynamics and structural biology to unveil the hotspots that differentially stabilize both states of RfaH. Our findings provide novel insights that will guide design of inhibitors blocking RfaH action.

## Introduction

Metamorphic proteins can access multiple structurally different and yet energetically stable states in solution (1), directly challenging the uniqueness of the native state considered in the thermodynamic hypothesis proposed by Anfinsen (2), typically interpreted as *one sequence - one fold*. This process takes place by major architectural rearrangements and is commonly related to changes in protein function and dynamics (3).

*Escherichia coli* RfaH is a metamorphic protein branching from the universally conserved NusG family of transcription elongation factors (4, 5), which enable processive RNA synthesis by RNA polymerase (RNAP) while simultaneously coupling it to concurrent processes (6). In NusG proteins, this coupling is achieved by two domains connected via a flexible linker. The N-terminal domain (NTD) is a structurally conserved α/β sandwich that freely binds the transcription elongation complex (TEC) by contacting the two largest RNAP subunits to form a processivity clamp around the DNA (7–9); the C-terminal domain (CTD) is commonly folded as a small five-stranded, antiparallel β-barrel able to interact with diverse cellular targets (10).

Despite sharing 41% sequence similarity, the structure of free *E. coli* RfaH displays striking differences from its paralog NusG. Instead of the canonical β-barrel, RfaH CTD is folded as an α-helical hairpin (αRfaH) which is tightly bound to the NTD, occluding the RNAP-binding site (Fig. 1) (11, 12). This autoinhibition is relieved upon domain dissociation, which is elicited during RfaH recruitment to the TEC or when the interdomain linker is cleaved, and the released CTD spontaneously refolds into the canonical β-barrel structure (βRfaH) observed in most NusG proteins (Fig. 1A) (12–14). This unique structural transformation is required to restrict RfaH action to just a few genes: autoinhibited RfaH is specifically recruited to the TEC in which an *ops* DNA sequence in the nontemplate DNA strand is surface-exposed (15, 16), most probably via an encounter complex (14); subsequently, domain dissociation leads to RfaH activation by CTD fold switching to attain a NusG-like structure (12, 13, 17) and by binding of the NTD to its high-affinity binding site on RNAP.

**FIGURE 1.**
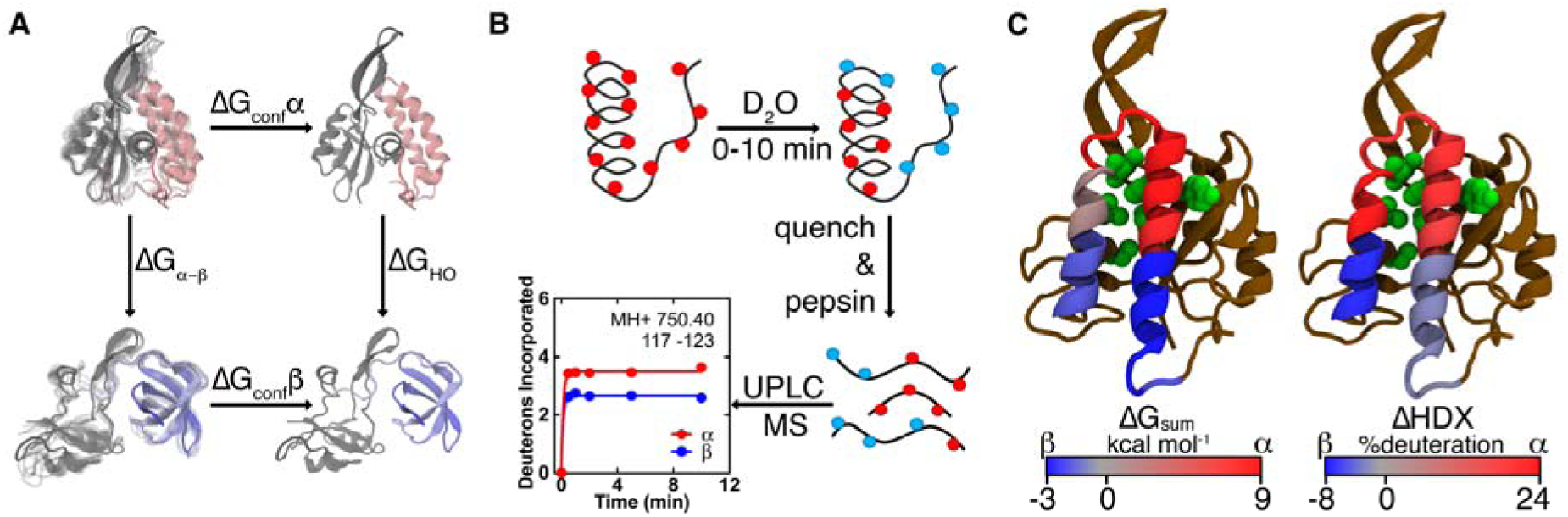
Computational and experimental assessment of local stability in the metamorphic protein RfaH. (*A*) Thermodynamic cycle of the confinement MD approach (25) used to estimate the per-residue ΔG between both RfaH folds. The autoinhibited form with the CTD in the α-state (pink, PDB 5OND) and the active form with the CTD in the refolded β-state that is canonical for NusG proteins (light blue, PDB 6C6S) are confined towards a deeply minimized state through a harmonic constraint (ΔG_conf_), allowing calculation of the difference in free energy between these structures (ΔG_HO_). (*B*) Scheme of the HDXMS experiments (26). Both full-length RfaH and the isolated CTD were incubated in deuterated buffer for different reaction times, quenched and pepsin-digested for analyzing the local extent of deuteron incorporation throughout RfaH. (*C*) Cartoon representations of full-length αRfaH, in which the CTD covers the RNAP-binding residues from the NTD (green), summarizing our findings from simulations (left) and experiments (right) on the differential local stability towards the α- (red) and β-state (blue) throughout the CTD of RfaH.

Since the trigger for RfaH metamorphosis is the complete *ops*-paused TEC (13, 14, 16), it is challenging to study this process experimentally. Instead, most of the thermodynamic and kinetic studies have used computational approaches to directly explore this fold-switch by simulating either the isolated CTD (18–20) or the entire RfaH protein (21, 22). Although the RfaH CTD is composed of only 51 residues (residues 112-162), its transition from alpha to beta has not been observed through conventional molecular dynamics (19), but through the use of enhanced sampling techniques (18, 20, 22) or reduced system granularity (21, 23). A way to circumvent such computing barriers is the use of confinement simulations, which rely on a discontinuous thermodynamic integration to estimate the absolute free energy of a clearly defined energy well (24). By evaluating two alternative states within the same system, one can calculate the energy required for the structural interconversion without explicitly observing such transition (25).

In this work, we employed hydrogen-deuterium exchange mass spectrometry (HDXMS), [^1^H,^15^N] heteronuclear single quantum coherence (HSQC)-based nuclear magnetic resonance (NMR) spectroscopy and confinement molecular dynamics (MD) to assess the differences in local stability between the autoinhibited and active folds of RfaH. By using deuterium as a probe to experimentally assess the solvent accessibility of peptides and individual backbone amides along RfaH in combination with simulations to estimate per-residue free energy changes upon refolding (Fig. 1), we aimed to trace back localized regions preferentially favoring the α- or β-fold and determine how this reversible switch is encoded within RfaH sequence.

## Materials and Methods

### Confinement Simulations

Structures of RfaH were built using the crystal structure of αRfaH (PDB 5OND) (16) and cryoEM composition of βRfaH (PDB 6C6S) (13). Rosetta3 suite was used to model and relax the flexible interdomain linker in both structures (27). To calculate the free-energy between both structures, implicit solvent MD simulations were performed on Amber16 along with CUDA as previously reported (28, 29). Although water molecules are not being directly computed, the per-residue root mean square fluctuations (rmsf) of RfaH in both structures is comparable to explicit solvent models previously reported (30–32). Furthermore, this method requires an initial step of deep energy minimization of the system, suggesting that the phase change the solvent would undergo during this process would result in more artifactual dynamics than those arising from a steady solvation potential. In total, 26 independent simulations were performed per system. For each, a harmonic position-restraining potential was used to drive the atoms towards a deeply energy-minimized configuration (see Supporting Material) for either the α- or β-state of RfaH. The stiffness of this potential was increased exponentially from mostly free (2.5·10^-5^ kcal mol^-1^ Å^-2^) to highly restrained (419.2 kcal mol^-1^ Å^-2^). Fluctuations and free energy were calculated for each basin as previously reported (Fig. S1) (28). Briefly, the estimated free energy is the result of a thermodynamic cycle (Fig. 1A) comprising two energy terms: confinement and convert. The former corresponds to the work applied by the external potential, exerted through all 26 simulations. Once confined, the magnitude of this work depends solely on the stiffness of the harmonic potential and not the configuration of the system (Fig. S1). Thus, beyond this point, the confinement free-energy difference between two distinct basins converges to a single value. The convert term corresponds to the free-energy difference between the already confined states, represented as the deeply energy-minimized configuration for each basin. This is calculated by using a harmonic oscillator approximation (28), in which the absolute free-energy of each state is calculated from the canonical partition function for a system of vibrating particles, whose frequencies are obtained from normal-mode analysis for both deeply minimized configurations (25). A per-residue free-energy decomposition scheme was also used as indicated previously (29). Formula derivations are described in the Supporting Material.

### Gene Expression and Protein Purification

All proteins sequences were encoded in plasmids harboring either a Tobacco Etch Virus protease cleaving site (TEV), or Thrombin cleaving site (HMK). Full-length *E. coli* RfaH was encoded into pIA777, a derivative of pET36b(+) containing its NTD–TEV–CTD–[His6] (33). The isolated CTD (i.e. RfaH residues 101 to 162) was harbored into a pETGB1A vector, containing [His_6_]-GB1-TEV-CTD (12). *E. coli* NusG was encoded into pIA244, a derivate of pET33 (34), encoding [His_6_]-HMK-NusG. Each plasmid was transformed onto *E. coli* BL21 DE3 expression system. Bacteria were grown at 37 °C until reaching an OD_600_ = 0.6-0.7, further induced with 0.2 mM IPTG (US Biological, Salem, MA, USA) at 30 °C overnight in the case of RfaH and its isolated CTD, or 30 °C for 3 hours for expressing NusG. The cells were further harvested by centrifugation at 4 °C.

Protein purification was performed by sonicating the cells at high intensity in buffer A containing 50 mM Tris-HCl pH 7.5, 150 mM NaCl and 10 mM imidazole pH 7.5. The protein-rich supernatant was obtained by centrifugation at 12,000 g for 30 min, further loaded onto a His-Trap HP column (GE Healthcare, Chicago, IL, USA), washed and then eluted in gradient with the same buffer supplemented with 250 mM imidazole pH 7.5. For isolated CTD, this eluate was incubated in buffer A with a non-cleavable His-tagged TEV protease at 4 °C overnight in a ratio of 20:1 mg of CTD:TEV protease. This mixture was then separated using another His-Trap HP column, collecting its flow-through enriched in isolated RfaH-CTD. Homogeneity of the protein purification was verified by SDS-PAGE.

Finally, all proteins were subjected to size exclusion chromatography prior to all experiments. This was performed on a Sephadex S75 column (GE Healthcare) connected onto an ÄKTA FPLC (GE Healthcare), using 20 mM Tris-HCl pH 7.9, 40 mM KCl, 5.0 mM MgCl_2_, 1.0 mM β-mercaptoethanol, 6.0% glycerol as the mobile phase.

### Hydrogen-Deuterium Exchange Mass Spectrometry

HDXMS was performed on each protein using a Synapt G2Si system with H/DX technology (Waters Corp, Milford, MA) as in previous works (35). In these experiments, 5 μL of protein solution at an initial concentration of 11 μM were allowed to exchange at 25 °C for 0-10 min in 55 μL of deuterated buffer containing 20 mM Tris-HCl pH 7.9, 40 mM KCl, 5.0 mM MgCl_2_, 1.0 mM β-mercaptoethanol, 6.0% glycerol. Then, reactions were quenched for 2 min at 1 °C using an equal volume of a solution containing 2 M GndHCl, 1% formic acid, pH 2.66. The quenched samples were injected onto a custom-built pepsin-agarose column (Thermo Fischer Scientific, Waltham, MA) and the resulting peptic peptides were separated by analytical chromatography at 1 °C. The analytes were electrosprayed into a Synapt G2-Si quadrupole time-of-flight (TOF) mass spectrometer (Waters) set to MS^E^-ESI+ mode for initial peptide identification and to Mobility-TOF-ESI+ mode to collect H/DX data. Deuterium uptake was determined by calculating the shift in the centroids of the mass envelopes for each peptide compared with the undeuterated controls, using the DynamX 3.0 software (Waters; Tables S1 and S2). The difference in deuteron incorporation of overlapping peptides was used for calculating the incorporation of overhanging regions when the difference in mass exceeded 5 times its uncertainty (see Supporting Material and Tables S3-S6). Incorporation was fitted to a single negative exponential per region (Fig. S2) to obtain the maximum deuteron incorporation per peptide, which was expressed as a percentage over the total number of amides. For formulation see Supporting Material, for the raw and processed data see Tables S3-S5.

### Hydrogen-Deuterium Exchange Heteronuclear Single Quantum Coherence Spectroscopy

^15^N-labeled full-length RfaH and RfaH CTD were produced as described (12). In brief, expression was carried out by growing *E. coli* in M9 minimal media (36, 37) supplemented with (^15^NH_4_)_2_SO_4_ (Campro Scientific, Berlin, Germany) as only nitrogen source. For the HDX experiments, the proteins were in 25 mM 4-(2-hydroxyethyl)-1-piperazineethanesulfonic acids (HEPES) pH 7.5, 60 mM NaCl, 5% (v/v) glycerol and 1 mM dithiothreitol (DTT), and spectra were recorded at 288 K for full-length RfaH and 298 K for the isolated CTD on Bruker *Avance* 700 MHz and *Avance* 800 MHz spectrometers, using cryo-cooled triple-resonance bearing pulse field-gradient capabilities. After lyophilization proteins were dissolved in D_2_O (99.98%) and the decay of signal intensities was observed in a series of [^1^H,^15^N]-HSQC spectra. Resonance assignment of backbone amide protons of RfaH were taken from a previous study (12). Exchange rates were determined by fitting the signal decay to a monoexponential curve (Table S6). The protection factors were calculated by dividing the experimental exchange rates (*k*_ex_) by the intrinsic exchange rates calculated from the amino acid sequence (*k*_rc_) and experimental conditions with tabulated parameters and were finally converted to ΔG values (Table S6) (38, 39).

### Data Availability

All experimental and computational data is available from the corresponding authors upon request.

## Results

### Confinement Molecular Dynamics Show Localized Differential Stability

To computationally ascertain the difference in local stability between both native states of RfaH, confinement MD simulations were used to estimate their global and local free energy differences. Given that interdomain interactions are critical for the stability of the CTD in the autoinhibited state (12), both structural states were modeled on the full-length RfaH to perform the confine-convert-release (CCR) approach (28, 29). These models were built in both states using the crystallographic αRfaH structure (corresponding to the autoinhibited state with NTD and CTD in the α-fold (16)) and the cryoEM βRfaH structure (corresponding to the activated, open state with NTD and CTD in the β-fold (13)), further refining the flexible loop connecting both domains using the knowledge-based Rosetta software (27). Then, these refined structures were used for confinement MD, thoroughly exploring the fluctuations from mostly free to highly restrained states, integrating and then decomposing the free energy difference between αRfaH and βRfaH required for such process (Fig. 2 and S1).

**FIGURE 2.**
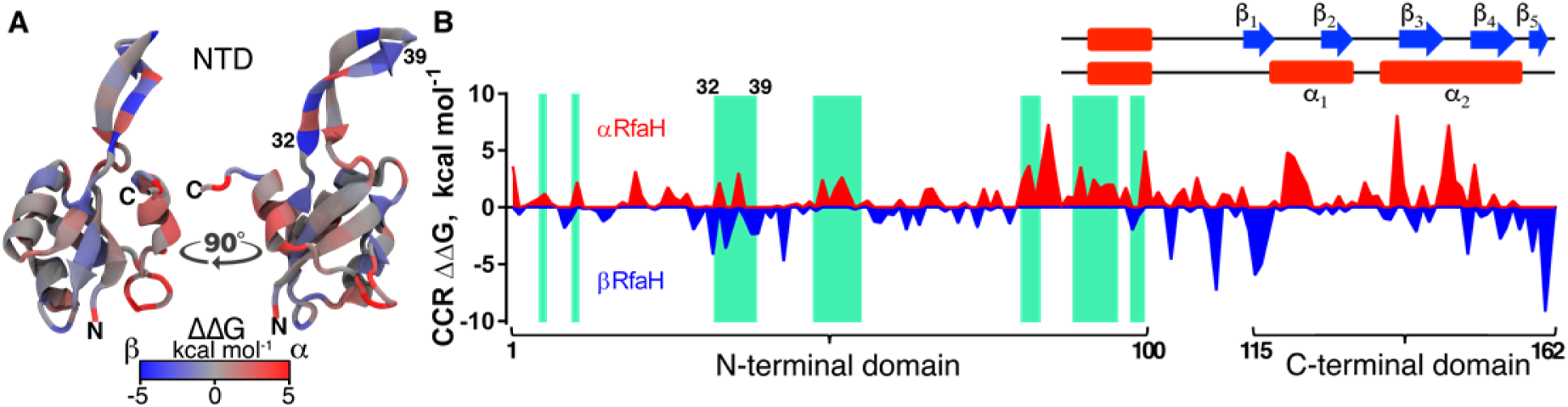
Confinement MD estimates anisotropic per-residue energetic contributions behind RfaH metamorphosis. Using the CCR approximation, the contribution towards stabilizing either αRfaH (red) or βRfaH (blue) metamorphic states was calculated (28, 29) and displayed onto the NTD structure (*A*) and along the RfaH sequence (*B*). The right-hand structure in *A* corresponds to a front view of the NTD interdomain interface (PDB 5OND). The green stripes in *B* highlight the NTD residues forming close contacts with the CTD in the crystal structure of αRfaH (16), with the β-hairpin residues 32-39 (also indicated in *A*) being the only NTD interface residues showing both stabilizing and destabilizing energetic contributions towards αRfaH. A visual guide for the fold-dependent secondary structure is shown for the CTD.

Free-energy decomposition into a per-residue level shows that localized groups of residues differentially stabilize either RfaH state. As shown experimentally (12, 40), most interdomain contacts deeply stabilize the autoinhibited αRfaH, with the exception of one region comprising the first strand in the β-hairpin of the NTD (residues 32-39 in Fig. 2) in which some residues are destabilizing. The behavior observed for these residues when compared to other interdomain NTD regions can be attributed to interactions between the CTD and the NTD β-hairpin observed in the structures of αRfaH (16) and βRfaH (13) used as starting configurations for the CCR approach. As this method drives the atom positions towards highly restrained states starting from these initial structures, the CTD-NTD interactions are stably kept for βRfaH throughout the confinement MD even though they are absent in solution (12). This can lead to overestimations of the contribution of these regions towards the stability of RfaH in the β-fold.

Although most of the CTD (residues 115-162) interacts with the NTD in αRfaH, per-residue free energy differences show localized heterogeneity in preferential stability towards each native state within this region. The C-terminal region of the linker (residues 110-114) and initial (residues 115-119) and terminal (residues 151-162) regions of the CTD show clear preference towards forming the strands β_1_ and β_4_-β_5_, respectively, whereas remaining residues 120-150 are more stabilized when forming the tip of the α-hairpin rather than strands β_2_ and β_3_ (Fig. 1C and 2). These results are consistent with previous simulations of the refolding pathway of RfaH in the context of the full-length protein using structure-based models (21). This is not the first example of heterogeneous and alternating ΔG along the primary structure using the CCR method; chameleonic proteins GA30 and GB30 provide per-residue free energies that strongly correlate to the sequence content from which they were engineered, effectively tracing back conformational space information from these simulations to their primary structure (29).

### Hydrogen/Deuterium Exchange Confirms Differential Stability of Metamorphic RfaH States

To determine how confinement calculations correlate with experimentally determined stabilities, HDXMS was performed with full-length RfaH, i.e. the CTD is in the α-fold, and isolated CTD, which is in the β-fold. This analysis identified a total of 43 different peptides derived from pepsin digestion of the 162 residues-long full-length RfaH (31 peptides for the NTD, 12 for the CTD), covering residues 7-159 (Fig. S2 and Tables S1 and S3). Meanwhile, 27 peptides were identified from pepsin digestion of the isolated CTD (residues 110-162), covering positions 117-162 (Fig. S2 and Tables S2 and S4). Given that many of the CTD peptides have varying lengths and overlapping regions between them, backbone amide deuteron incorporation was deconvoluted into 7 unique regions that were observable for RfaH CTD in both the α-fold and the β-fold (see Supporting Material) and covered almost the entirety of the CTD (residues 117-159).

As measure for flexibility, we determined the relative deuterium uptake of full-length RfaH, i.e. αRfaH, and the isolated CTD, i.e. βCTD (Fig. 3), corresponding to the ratio between the maximum deuterium incorporation, calculated as the saturation value of an exponential fit to the deuterium uptake (Fig. S2), and the maximum theoretical incorporation, which depends on the peptic fragment sequence and length (41). In the NTD, buried regions show incorporation of about 30%, while solvent-exposed regions exhibit increased deuteration of around 50% (Fig. 3A and S2). Strikingly, in full-length RfaH almost all of the peptides of the CTD display between 40-50% deuteron incorporation, with the exception of a single region covering residues 130-142, whose incorporation reaches a maximum of only 23% (Fig. 3B). This is reminiscent of the temperature factors observed in the crystal structure of full-length RfaH, wherein residues 115-128 and 153-156 display B-factor values over 50 while residues covering the tip of the α-hairpin display values of around 30 (16). Thus, these results indicate that the ends of the α-hairpin are highly flexible whereas the tip is relatively more rigid.

**FIGURE 3.**
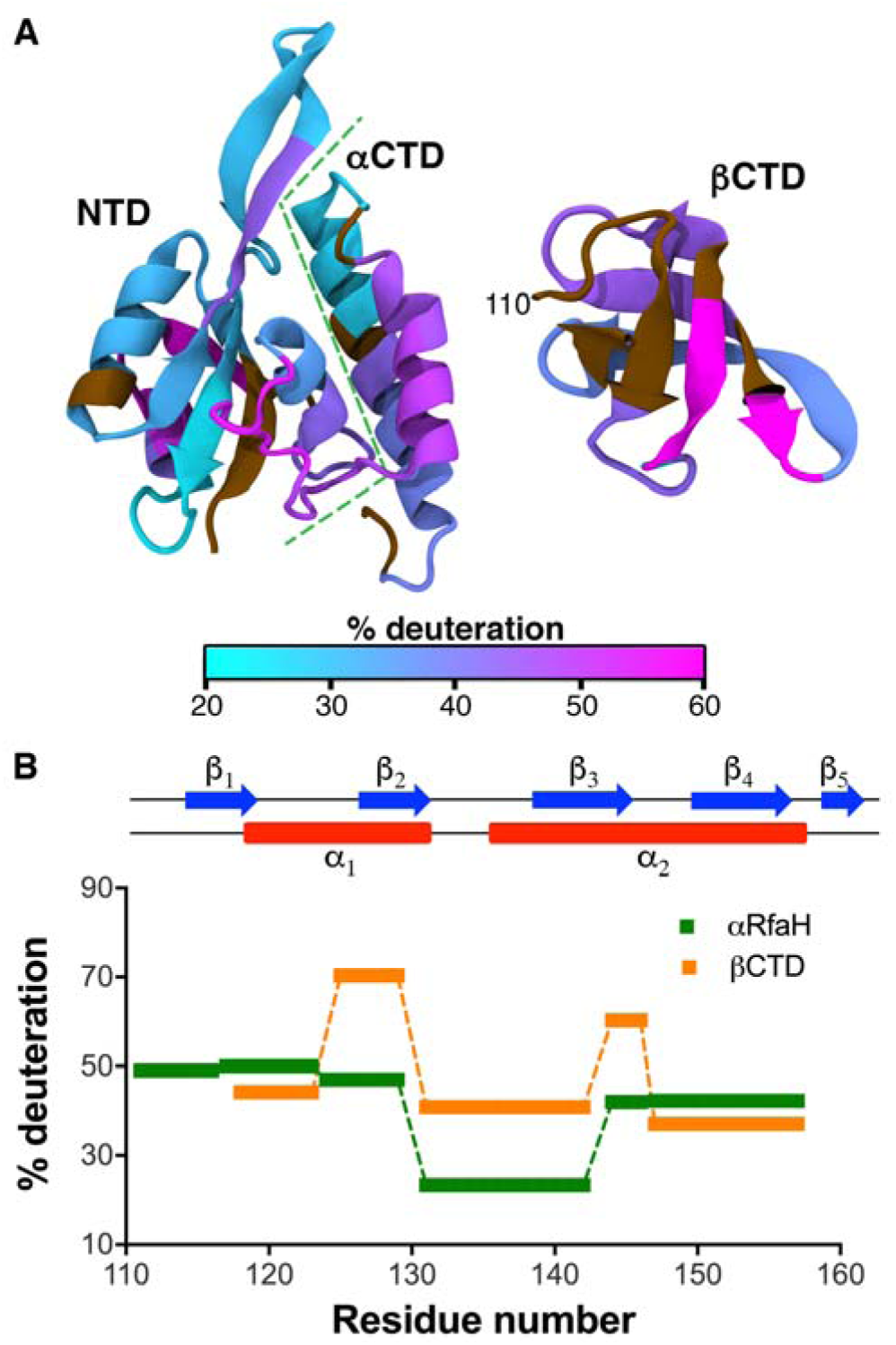
Mapping of the differences in local flexibility between RfaH states by HDXMS. (*A*) Structural mapping of the relative deuterium uptake of different regions of RfaH NTD and CTD in the context of the full-length RfaH (αRfaH, PDB 5OND) or the CTD in isolation (βCTD, PDB 2LCL). Regions are colored with a gradient from cyan (solvent-protected against deuterium exchange) to magenta (solvent-accessible). Residues whose deuteron incorporation could not be determined by mass spectrometry are shown in brown. (*B*) Relative deuterium uptake for RfaH CTD in its α-and β-state (green and orange, respectively). Each rectangle represents a given peptide whose extent of deuteron incorporation was determined by mass spectrometry.

In contrast to αRfaH, most regions of the isolated CTD display around 40% relative deuterium uptake with the exception of a short loop (residues 144-146) and strand β_2_, which exhibit deuteration extents of around 60 and 70%, respectively. For comparison, the experiment was also carried out with the full-length NusG protein (Fig. S3 and Table S5), whose CTD is always folded as a β-barrel that does not stably interact with its NTD (10). Relative deuterium uptake between 50-60% is observed for almost all regions within NusG CTD, slightly higher than those observed for its metamorphic paralog, except strand β_2_ that exhibits 30% less exchange as compared to RfaH. Despite having superimposable structures and 43% sequence similarity, RfaH CTD residues have overall larger aliphatic and more hydrophobic side chains than those of NusG, physicochemical features that are compatible with the observed lower flexibility of RfaH CTD in the β-fold. This analysis shows that, nevertheless, RfaH CTD dynamics in the β-fold are not drastically different from that of its paralog NusG, while highlighting the strongly reduced flexibility of the tip of the α-helical hairpin in αRfaH.

To further confirm the heterogeneity in local stability and flexibility observed for RfaH, we performed NMR-based H/D exchange experiments on full-length RfaH and the isolated CTD. The lyophilized ^1^H,^15^N-labeled proteins were dissolved in D_2_O and HDX was monitored via the decay of signal intensities in a series of two dimensional [^1^H,^15^N]-HSQC spectra. In full-length RfaH only 17 signals in the NTD and 5 in the α-folded CTD corresponding to individual amides were detectable and analyzed (Fig. S4 and Table S6). All other amide protons in full-length RfaH and all amide protons in the isolated CTD exchanged too fast (i.e. the exchange was complete within the experimental time for the first spectrum), thus suggesting that these amides are either solvent-exposed or not stably involved in hydrogen bonds to observe them in NMR-based HD exchange.

The decay rates in signal intensity of the observable amides were fitted onto a single exponential and converted into exchange protection factors (PFs), which equate to an equilibrium measurement of local stabilization in a folded conformation as compared to the unfolded state and can be further used to determine the free-energy change involved in exposing the protein amides to the solvent (Table S6) (38, 39). Remarkably, all of the 22 analyzable RfaH NTD and CTD amide protons are located in regions of preferential stability towards the α-fold according to CCR (Fig. S4), with all the CTD signals located on the tip of the α-helical hairpin. Also, these single amides are encompassed in peptides showing low deuteration in HDXMS experiments using full-length RfaH (Fig. S4). Thus, these data confirm that the tip of the α-helical hairpin exhibits a stability comparable to that of the NTD in full-length RfaH.

To provide greater detail into the similarities in local stability of RfaH CTD in the α-fold and the β-fold observed through HDXMS and MD, the ΔG values were summed for residues matching the 5 regions experimentally observed through HDXMS. Comparison of the local differential stability patterns using both strategies revealed striking similarities in distribution as well as in magnitude (Fig. 4), with the exception of CTD residues 124-129 from helix α1 that constitute part of the tip of the α-helical hairpin (Fig. 1). This is partly explained by the high solvent accessibility of this region in the β-barrel fold as ascertained by HDXMS, exhibiting the highest extent of deuteration (Fig. 3). In contrast, the confinement procedure of the MD simulations might stabilize the interactions within this region, thus allowing their energetics to be comparable to those estimated for the CTD in the α-fold (Fig. 2).

**FIGURE 4.**
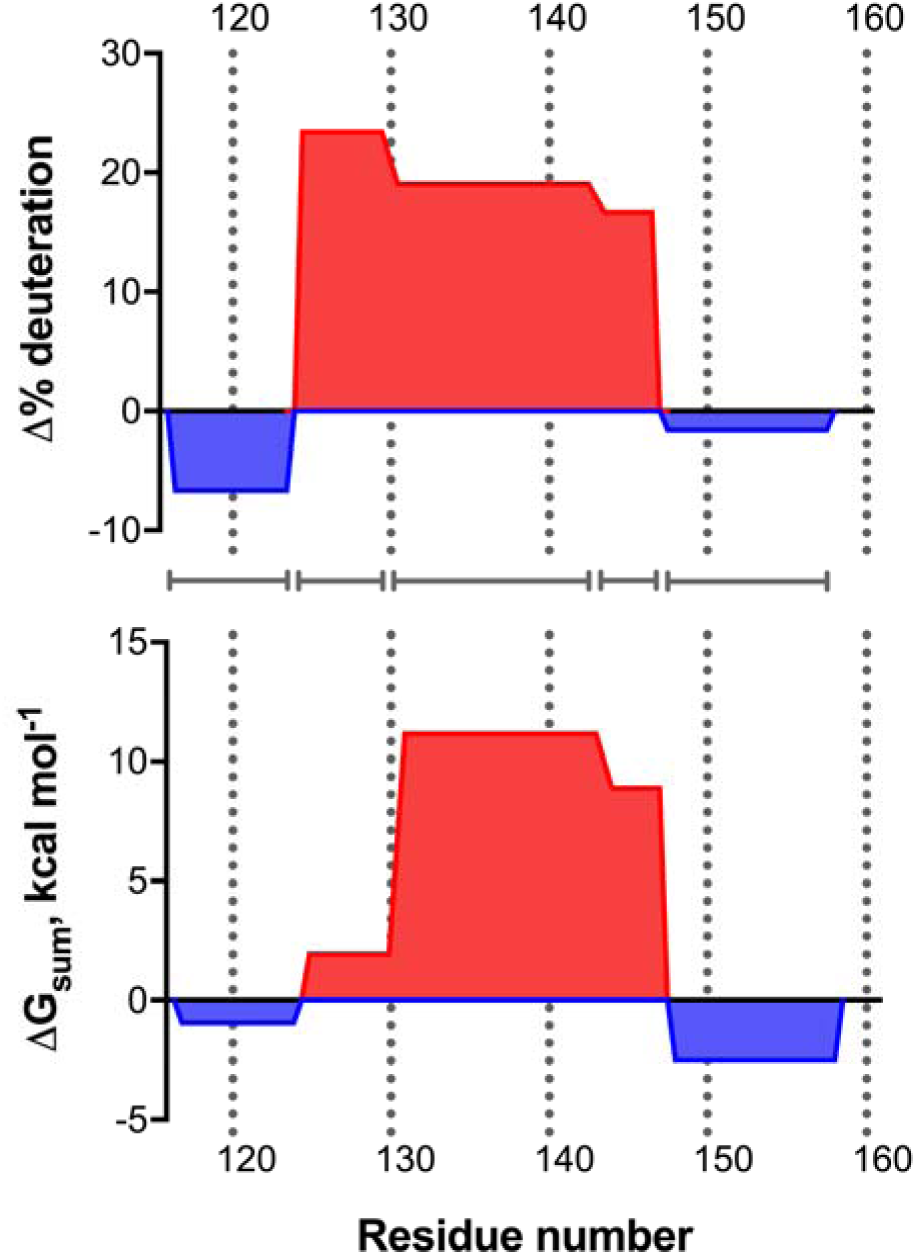
Comparison between computationally-derived free energy (ΔG_sum_) and experimentally-determined flexibility (Δ% deuteration) between 5 different regions of RfaH CTD (indicated by grey lines) in its α-fold and β-fold. Per-residue free energy differences were added accordingly to match the length of the peptides analyzed via HDXMS. Regions in red and blue have preferential stability towards the α-fold and β-fold, respectively.

## Discussion

The molecular mechanism by which the transformer protein RfaH completely refolds its CTD during its activation has been experimentally elusive. Computational and experimental approaches strongly support the importance of interdomain contacts in controlling RfaH metamorphosis (12, 21, 40, 42). In this work, both hydrogen-deuterium exchange and confinement MD reveal that the interaction-rich upper region of the α-helical hairpin, comprising residues 125-145, provides the highest degree of stabilization towards the α-folded CTD (Fig. 1C and 4). Moreover, confinement MD and [^1^H,^15^N]-HSQC-based HDX experiments show that interdomain contacts stabilize to a similar extent the tip of the α-helical CTD as well as the NTD (Fig. S4). This suggests that the structural metamorphosis of RfaH from the autoinhibited to the active state is controlled not only by native contacts with its binding partner, but also by intrinsic CTD determinants within the aforementioned region. Thus, it comes as no surprise that the computational analysis of NusG and RfaH sequences identified at least seven residues that are highly conserved within the RfaH subfamily and significantly contribute to NTD-CTD binding, of which three NTD (E48, F51 and P52; *E. coli* numbering) and three CTD residues (F130, R138 and L142) are located in the vicinity of the α-helical hairpin (12, 42).

Using structure-based models, we previously simulated RNAP binding to RfaH, and concluded that its binding in the vicinity of NTD residue E48, adjacent to the tip of the hairpin, may suffice to favor refolding of the CTD towards the β-fold, and thus the relief of autoinhibition (21). Our present results, along with the recent cryo-EM structure of RfaH bound to the complete TEC (13), are consistent with this hypothesis. Binding of the autoinhibited αRfaH, which cannot contact its high-affinity site on the β’ subunit of RNAP, is thought to be mediated by its initial contacts to a hairpin that forms in the non-template DNA strand and to the β subunit gate loop (13, 14, 16, 43), resulting in an encounter complex (14).

The preference towards the β-fold displayed by residues 110-119 and 151-162 (Fig. 2) strongly suggests that refolding of the CTD towards the β-fold starts with the unfolding of the ends of the α-helical hairpin, as they fluctuate towards a locally unfolded state even before dissociation (Fig. 3). Thus, the tip of the α-helical CTD acts as an anchor preventing its spontaneous refolding into a β-barrel. This is supported by previous observations that disruption of the E48:R138 salt bridge located in this region led RfaH to exist in equilibrium between the autoinhibited and active folds (12). However, this view contrasts with conclusions drawn from other computational approaches (18, 22), which suggested that contacts involving RfaH CTD residues comprising the strand β_3_ are particularly stable and nucleate the β-barrel, as solution dynamics do not display high stability in this region for RfaH or NusG CTD in the β-fold (Fig. S3).

Folding of RfaH into a stable, autoinhibited structure is essential for its function. Since RfaH has a higher affinity for the TEC than NusG (13), its binding to RNAP has to be tightly controlled to prevent misregulation of NusG-dependent housekeeping genes. The emergence of the autoinhibited state, which is relieved only in the presence of a 12-bp *ops* DNA element with complex properties (13, 16), presents an elegant solution to this problem. Our results suggest that establishing interdomain contacts at the tip of the hairpin, blocking most of the RNAP-binding residues (Fig. 1C), is sufficient to enable this autoinhibition. Moreover, the emergence of this novel fold causes only few changes in the local stability and dynamics of the canonical β-barrel of NusG CTDs (Fig. 3 and S3), supporting its ability to interact with the translational machinery (12, 14). These interactions, established with the ribosomal protein S10, are formed through hydrophobic residues located in strands β_2_ (residues 122-126) and β_4_ (residues 145-148) (6, 12, 14) whose identities are mostly conserved between RfaH and NusG. Our results show that these residues are stably interfaced with the NTD, thus explaining why S10 is unable to elicit RfaH metamorphosis on its own (12).

The key differences between RfaH and NusG, metamorphosis and sequence divergence of the CTD, underpin their orthogonal cellular functions. Even though NusG and RfaH bind to the same site on the TEC and display similar effects on RNA synthesis (5, 13), they paradoxically play opposite roles in the expression of horizontally acquired genes. NusG cooperates with Rho to silence foreign DNA, an activity that explains the essentiality of *E. coli* NusG (44), through direct contacts between NusG-CTD and Rho (45). In contrast, RfaH does not interact with Rho and abolishes Rho-mediated termination in its target operons, all of which have foreign origin, in part by excluding NusG (43). Remarkably, grafting a five-residue NusG loop (S^163^IFGR) into RfaH (N^144^LINK) creates an even more potent activator of Rho (45). Thus, the loss of interactions with Rho, another pivotal step in the evolution of RfaH, also occurs around the tip of the helical hairpin.

Altogether, our results show that the tip of the α-helical hairpin is the main determinant for stabilizing the autoinhibited state of RfaH, and that this localized stability arises from interdomain interactions and intrinsic sequence-encoded preferences. Both our present results and other available evidence suggest that targeted substitutions in this CTD region enabled both the acquisition of the autoinhibited state, in which this region forms the tip of the stabilizing α-hairpin, and the loss of termination-promoting contacts with Rho. These changes converted a nascent paralog of NusG, an essential xenogene silencer, into an activator of horizontally transferred virulence genes that encode capsules, toxins, and conjugation pili (46). We hypothesize that the molecular details about RfaH mechanism can be harnessed to design ligands that interfere specifically with RfaH activity and thus virulence (47). In addition to directly inhibiting the expression of virulence genes, these ligands may also limit the spread of plasmid-encoded antibiotic-resistance determinants through conjugation and synergize with the existing drugs by compromising the cell wall integrity in Gram-negative pathogens.

## Acknowledgements

This research was funded by Fondo Nacional de Desarrollo Científico y Tecnológico (FONDECYT 11140601) and International Cooperation Grant (REDI170624) from Comisión Nacional de Investigación Científica y Tecnológica (CONICYT), the National Institutes of Health (NIH 1S10OD016234), the NVIDIA Corporation through the GPU Grant Program, and the German Research Foundation (Ro 617/21-1). CARS was funded by ASBMB, PABMB and IUBMB through the PROLAB program. PGD and JAM were funded by CONICYT Doctoral Scholarships (CONICYT-PFCHA 21181705 and 21181787, respectively). We acknowledge the gracious help of Dr. Ken Dill’s group at Stony Brook University and Dr. Arijit Roy regarding the use and calculation of the CCR approach used in this work. We finally thank Prof. Paul Rösch and Dr. Kristian Schweimer for helpful discussions on the manuscript.

## Author Contributions

P.G.D., E.K., I.A. and C.A.R.S. designed the research. J.A.M., S.S., S.H.K. and C.A.R.S. conducted the experiments. P.G.D., E.K., S.H.K. and C.A.R.S. analyzed the data. P.G.D. and C.A.R.S. conducted and analyzed the computational work. P.G.D., E.K., S.H.K., I.A. and C.A.R.S. wrote the manuscript.

## Supporting Citations

References (48-52) appear in the Supporting Material.

## Supporting Material

Supporting Methods, Figures S1-S4 and Tables S1-S6 appear in the Supporting Material.

